# Parchment Glutamine Index (PQI): A novel method to estimate glutamine deamidation levels in parchment collagen obtained from low-quality MALDI-TOF data

**DOI:** 10.1101/2022.03.13.483627

**Authors:** Bharath Nair, Ismael Rodríguez Palomo, Bo Markussen, Carsten Wiuf, Sarah Fiddyment, Matthew James Collins

**Affiliations:** Globe Institute, Faculty of Health and Medical Sciences, University of Copenhagen – Copenhagen, Denmark; McDonald Institute for Archaeological Research, University of Cambridge – Cambridge, United Kingdom; Department of Mathematical Sciences, University of Copenhagen – Copenhagen, Denmark

**Keywords:** ZooMS, MALDI-TOF, biocodicology, deamidation, collagen, parchment, PQI

## Abstract

Parchment was used as a writing material in the Middle Ages and was made using animal skins by liming them with Ca(OH)_2_. During liming, collagen peptides containing Glutamine (Q) undergo deamidation resulting in a mass shift of 0.984 Da. Assessing the extent of deamidation can inform us about parchment production patterns and quality. In this study, we propose a simple three-step workflow, developed as an R package called `MALDIpqi()`, to estimate deamidation in parchment derived collagen using low-resolution MALDI-TOF spectra. After pre-processing raw spectra, we used weighted least-squares linear regression to estimate Q deamidation levels from the convoluted isotopic envelope for seven collagen-peptide markers. Finally, we employed a linear mixed effect model to predict the overall deamidation level of a parchment sample termed Parchment Glutamine Index (PQI). To test the robustness of the workflow, we applied `MALDIpqi()` to previously published ZooMS data generated from almost an entire library of the Cistercian monastery at Orval Abbey, Belgium. In addition to reliably predicting PQI, we observed interesting patterns pertaining to parchment production. `MALDIpqi()` holds excellent potential for biocodicological and other archaeological studies involving collagen, such as bone, but we also foresee its application in the food and biomedical industry.

## Introduction

Glutaminyl and asparaginyl residues are molecular clocks which deamidate with predetermined half-lives (NE Robinson, Zabrouskov, et al., 2006). Glutamine (Gln, Q) deamidation occurs via two mechanisms, 1) direct hydrolysis, and 2) through the formation of a cyclic imide intermediate. Irrespective of the mechanism, instability of reaction intermediates results in slower deamidation rates of Gln. Because the deamidation rates of glutaminyl residues are slower than asparaginyl residues (NE Robinson, ZW Robinson, et al., 2004; Tonie Wright and Urry, 1991), it has been advanced that Gln deamidation could be a better tool at our disposal to investigate chemical processes such as assessing the quality of skins in the food and leather industries (Maffia et al., 2004) and the age of fossils (Doorn et al., 2012), although in the latter case (Schroeter and Cleland, 2016) argue that Gln deamidation is an indicator of preservational quality and environmental conditions rather than age (and authenticity) of ancient proteins. Here, we use Gln deamidation to assess variability in parchment production.

Mass spectrometry is well suited to detecting sites of deamidation, which increases the molecular mass of deamidated peptides by 0.984 Dalton (Da), and is easily detected and localised in the sequence by a mass shift in the tandem mass spectra. Since deamidation results in a mass shift of one nominal Da higher, the isotopic envelopes of deamidated and non-deamidated peptides overlap each other in MS1 spectra resulting from low-resolution mass spectrometers.

We explore a mathematical approach to derive the level of deamidation in glutamine from matrix-assisted laser desorption/ionisation time-of-flight (MALDI-TOF) MS1 spectra. MALDI-TOF mass spectrometry is widely applied in bioarchaeology for species identification using peptide mass fingerprinting (PMF) of collagen, known as ZooMS(Buckley, Collins, et al., 2009; Buckley, Whitcher Kansa, et al., 2010). Herein, we use an isotopic envelope deconvolution method (similar to the approach used by (Wilson et al., 2012)) to estimate the extent of glutamine deamidation in selected tryptic peptides but then integrate the individual deamidation estimates to derive an overall index for a given sample. In order to develop the method we have used published MALDI spectra of parchment (e.g. (Fiddyment, Holsinger, et al., 2015)) which we have then applied to a newly released data set from Orval Abbey (Ruffini-Ronzani et al., 2021).

Parchment is the dehaired and limed skin of an animal (Reed, 1972; Ryder, 1964). Liming is typically the first stage in parchment and leather preparation;it loosens the hairs from the hides, swells the collagen and saponifies some of the skin lipids prior. Gln deamidation occurs when the skins are soaked in lime, a solution of calcium hydroxide Ca(OH)_2_, which at ambient temperature has an average pH of about 12.4 and is used in different strengths during the parchment making process. The alkaline environment results in direct side chain hydrolysis of the amide group on asparagines and glutamines. A longer exposure (or higher concentration and/or temperature) of lime results in an increase in the extent of deamidation. If not controlled correctly, an excessive exposure to lime can compromise the integrity of skin and to weaken it to such a degree that is no longer usable. By measuring the level of deamidation present in different samples we can start to assess the different production qualities from different regions and time and correlate this to prices and availability of parchment obtained from historic records. Consequently, the extent to which these skins are limed can be interrogated through the measurement of the level of glutamine deamidation.

By assessing the relative rates of deamidation of different tryptic peptides we derived a single value (with associated errors) which we term the Parchment Glutamine Index (PQI). Samples which retain the most intact glutaminyl residues have the highest PQI values; as deamidation increases, PQI falls. In MALDI-TOF mass spectrometry baseline noise in the spectra (Kolibal and Howard, 2006; Krutchinsky and Chait, 2002) results in a distortion of the relative intensity of the peaks across an isotope envelope which in turn affects estimates of deamidation based on the deconvolution of the envelope. Consequently values greater than 1 (ie. no deamidation) are possible due to noisy baselines, while values close to 0 are never observed in parchment this would follow complete gelatinisation.

## Materials

In order to establish the model, we used available published ZooMS data to establish correlations between the rates of deamidation of different tryptic peptides. We then test our model using data generated from almost the entire library of the Cisterian monastery at Orval Abbey, Belgium (Ruffini-Ronzani et al., 2021). Explanation of the data generated can be found in the data article (Bethencourt et al., 2022) and the ZooMS data is uploaded in Zenodo (https://doi.org/10.5281/zenodo.5648106).

## Methods

### Sampling and spectra acquisition

Parchment sampling was done by a non-invasive triboelectric extraction of collagen following a previously published method (Fiddyment, Holsinger, et al., 2015) wherein non-written areas of parchment surface was gently rubbed with an eraser followed by the collection of crumbs in a 1.5 mL Eppendorf tube using acid free paper. Freshly cut piece of polyvinylchloride (PVC) eraser was used for each parchment sample and nitrile gloves were worn during the procedure.

Samples collected in Eppendorf tubes were added with 75 *μL* of 50 mM ammonium bicarbonate buffer (pH 8) and 1 μL of trypsin (0.4 μg μL^-1^) and incubated at 37 °C for 4 hours to digest collagen. Eppendorf tubes were then spun-down on a benchtop centrifuge at maximum speed and enzyme digestion was quenched by adding 1 μL 5%*v/v* trifluoroacetic acid (TFA). After transferring the supernatant to fresh Eppendorf tubes, collagen peptides were extracted using C18 resin ZipTip (add round R) pipette tips and eluted into 50 μL of 50 % acetonitrile (ACN) and 0.1 % TFA. Spotting (1 μL) was done in triplicates on a ground steel plate and mixed with 1 μL of matrix solution (α-cyano-hydroxycinnamic acid), along with a calibration mixture (Fiddyment, Holsinger, et al., 2015; Fiddyment, Teasdale, et al., 2019).

Samples were analysed by MALDI-TOF mass spectrometry at BioArch laboratory at the University of York using a Bruker Ultraflex III mass spectrometer with a mass range of 800-4000 *m/z* at the Center of Excellence in mass spectrometry.

### Selection of peptides

In order to assess the overall PQI we used as many peptides as possible and selected them based upon the following criteria:

1. They contain at least one glutamine.
2. They are consistently and reliably detected in the MALDI-TOF MS analysis.
3. They are present in all three species used to make parchment (calf, sheep and goat). All these peptides have the same mass in the different species, except *m/z* 3033 and 3093, which are the equivalent peptides for calf/sheep and goat, respectively.

A final list consisting of eight peptides was compiled (Table 1), of which a maximum of seven can be detected in any one sample due to the equivalence of peptides *m/z* 3033 and 3093), and used to run the subsequent analysis.

**Table 1.**
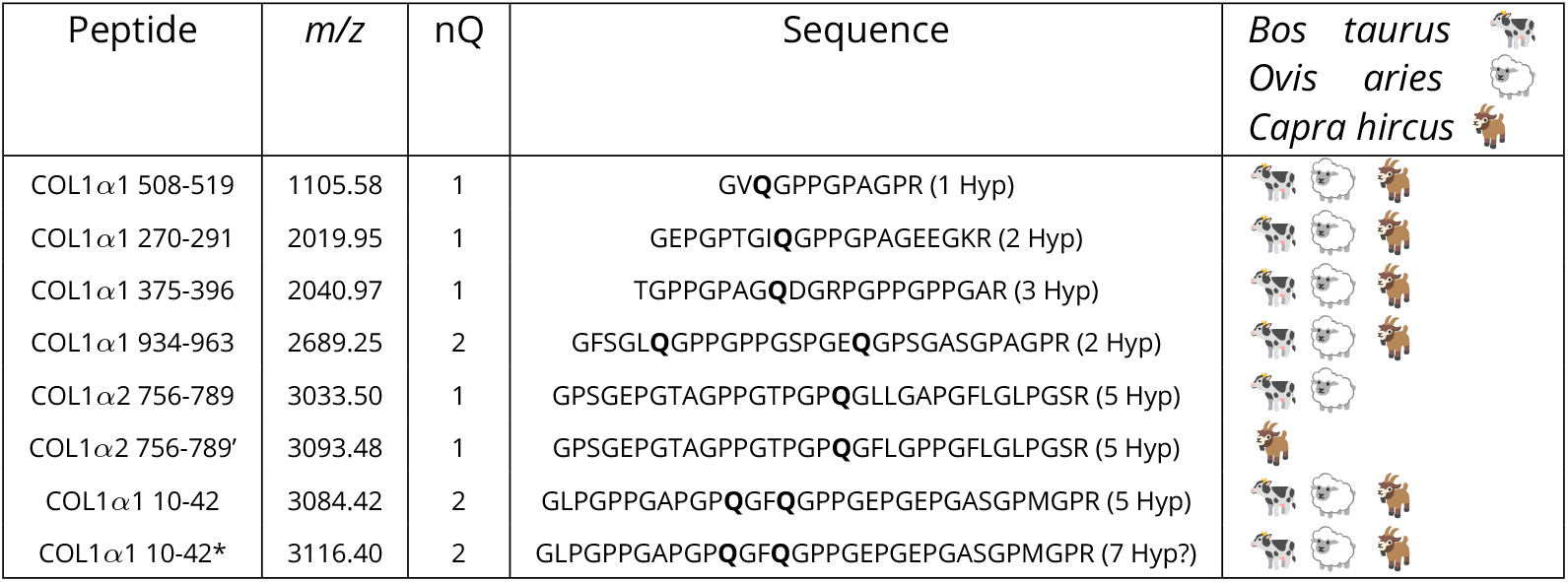
List of peptides used in the analysis, using the (Brown et al., 2021) nomenclature for ZooMS marker peptides. Sequences for each mass were inferred from Mascot analysis of parchment datasets (SF and JW personal communication). All masses are consistent with typical hydroxylation patterns of collagen except where indicated by ”?”. To differentiate the peptide COL1*α*2 756-789 from cow and sheep from the goat one, we have marked the latter with ”‘”. Similarly, the peptide COL1*α*1 10-42 with 7 hydroxyprolines is marked with “*”

### Pre-processing of raw data

We performed pre-processing of the spectra using the R (R Core Team, 2021) package MALDIquant (Gibb and Strimmer, 2012):

- The Savitzky-Golay-filter (Savitzky and Golay, 1964) smoothed the spectra and reduced small, highly frequent noise. This allows for better subsequent baseline and noise estimation and peak maxima determination. We used a moving half-window size of 8, following the recommendation by (Bromba and Ziegler, 1981) of keeping it smaller than the full width at half maximum of the peaks.
- We estimated the baseline (and then subtracted) using the Statistics-sensitive Non-linear Iterative Peak-clipping algorithm (SNIP) (Ryan et al., 1988) implemented in MALDIquant; the iterations parameter of the algorithm is set to 20.
- We estimated noise using the SuperSmoother (Friedman, 1984) method. Peaks are detected if they are a maximum with a half-window size of 20 and are above a signal to noise ratio (S/N) of 1.5.
- Finally, we extracted isotopic-like distributions for each of the selected peptides by finding the canonical *m/z* value and 5 following peaks (if detected) at the isotopic distance of 1 Da, allowing for a small tolerance deviation of 1.5 ·10^-4^·*mass* units. We discarded incomplete isotopic distributions in the peptide spectra when, 1) a peak other than the first or the last or just the first one is missing, or 2) less than four peaks are identified.

Figure 1 shows the spectra before and after pre-processing for five selected samples.

**Figure 1.**
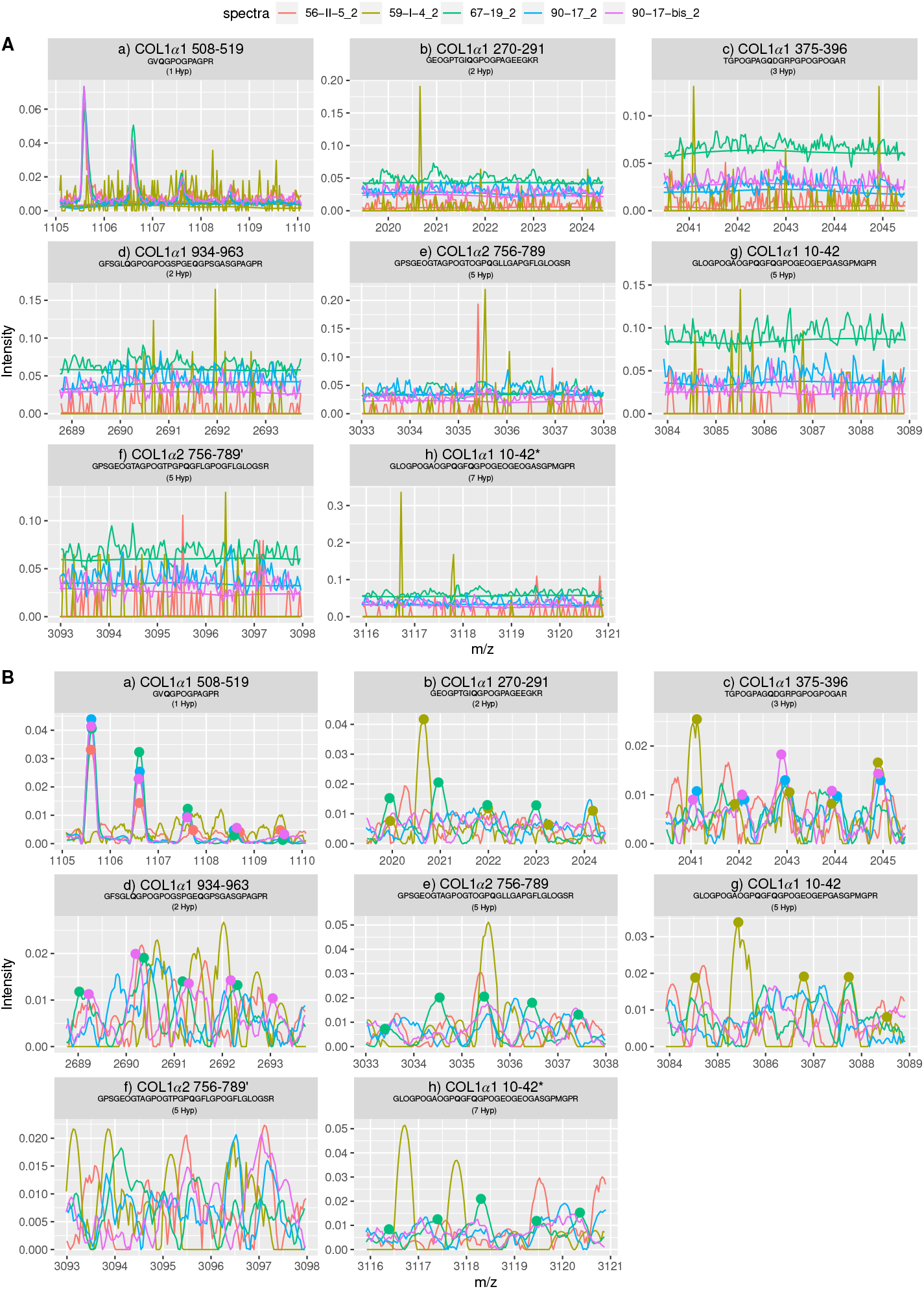
Illustration of overlapping isotopic distribution (up to 6 peaks) of deamidated and non-deamidated fraction of peptides used in the model before (A), and after preprocessing (B). We show the baseline estimated on the smoothened spectra and the peaks that are detected after the preprocessing step 3). Note the variation in intensity and the difference in the S/N of peaks for each peptide. Whereas there is a clear distinction of individual peaks (hence, high S/N) for COL1*α*1 508-519, peak distinction becomes complex for COL1*α*1 10-42 or COL1α2 756-789 due to the noisy spectra (hence, less S/N). Five randomly selected samples are shown here.

### Estimation of deamidation level of peptides

Deamidation of a peptide consisting of glutamine (Q) at a single site results in a mass shift of approximately +0.984 Da so that the first peak of the isotope distribution for the deamidated peptide coincides with the second peak of the isotope distribution for the non-deamidated peptide (at the resolution of our data). For a peptide with *k* possible deamidation sites, each additional deamidation results in a further +0.984 Da mass shift leading to k overlapping isotope distributions. The level of deamidation of a peptide can be estimated by deconvoluting the two overlapping isotopic distributions.

#### Theory

In order to explain the method, we focus on one peptide and assume there are m + 1 isotopic peaks of the peptide available with isotope distribution

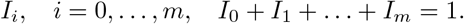

We followed the method as described in (Wilson et al., 2012) to calculate theoretical isotopic distributions for the peptides. For convenience, we put *I_i_* = 0 for *i* < 0. During deamidation, we expect a shift in the isotope distribution

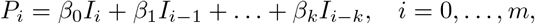

where 1 ≤ *k* < *m* and *β_ℓ_* ≥ 0, *ℓ* = 0,…, *k*, is the probability that *ℓ* positions are deamidated, such that *β*_0_ + *β*_1_ +…+ *β_k_* = 1. It holds that *P*_0_ + *P*_1_ +…+ *P_m_* = 1. In the current study, we take *k* =1.

#### Weighted least square and linear regression

We developed a general theory assuming *m* +1 measurements, one for each isotope, replicated n times. However, to estimate the overall deamidation level from multiple peptides and replicates simultaneously (see Section) we obtained estimates of the deamidation level for each of the 3 replicates separately, that is, we apply the theory below with *n* =1.

Notation for observed intensities of each isotopic peak and replicate:

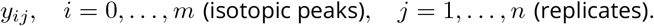

There might be missing values and/or missing replicates.

The measurements are proportional to *P_i_*, *i* = 0,…, *m*, hence in particular the measurements do not sum to one. In general, consider the linear model

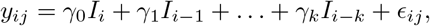

where *γ_ℓ_* ≥ 0, *ℓ* = 0,…, *k*, are parameters, and *ϵ_ij_* is (unobserved) noise. The deamidation fractions are obtained as

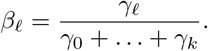

We avoided assuming a specific noise structure (for example, normal distributed noise) and used weighted least square to estimate the unknown parameters,

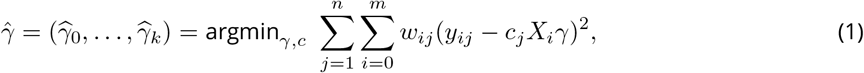

where *γ* = (*γ*_0_,…, *γ_k_*)^*T*^ is a column vector and *c* = (*c*_1_,…, *c_n_*) is a row vector. Here *c_j_* is a scaling factor for the *j*’s replicate with *c*_1_ = 1. The idea being that replicates show the same trend but might vary in signal intensity, hence scaling is required to adjust the parameters. Furthermore, *w_ij_* is a weight for the *y_ij_*’s data point, and *X_i_* is the *i*th row of

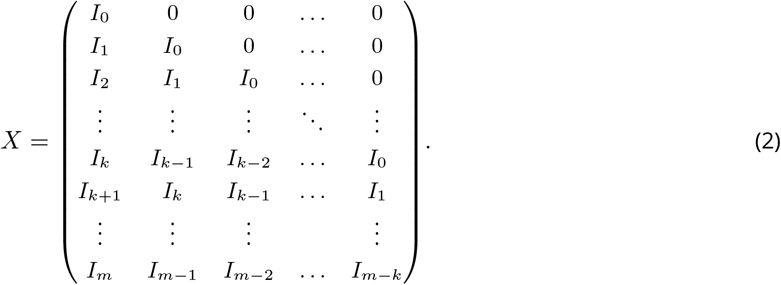

In total there are (*k* + 1) + (*n* – 1) = *k* + *n* parameters and *n*(*m* + 1) measurements, assuming none are missing. If measurements are missing the corresponding terms in Equation (1) are omitted.

The estimates can be obtained by weighted linear regression with design matrix *X* and diagonal weight matrix

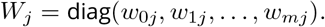

Let 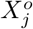 be the matrix *X* with the rows corresponding to the missing measurements of replicate *j* omitted (upper index *o* for omitted), let 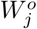 be the matrix *W_j_* with the rows and columns corresponding to the missing measurements of replicate *j* omitted, and let 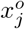 be the column vector with missing measurements or replicate *j* omitted.

Then the estimates can be obtained iteratively by

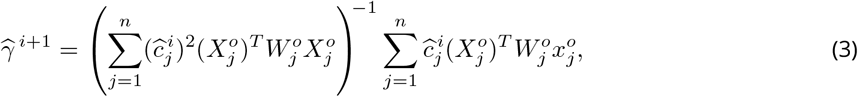

and

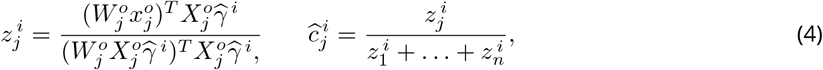

where 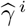 and 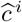 are the ith iterated estimates of the column vectors *γ* and *c* = (*c*_1_,…, *c_n_*). One continues until the difference between consecutive estimates is small with initial estimate 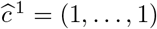.

Equation (3) is the standard weighted least square estimate assuming the scaling factors are known. Equation (4) is an update of the scaling factors assuming the other parameters are known.

For use in the later stage of the workflow to estimate the overall deamidation level (see Section) we define the *Reliability measure*

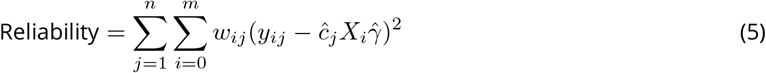

If there is only one replicate or one estimates *γ* separately for each replicate (*n* = 1), then

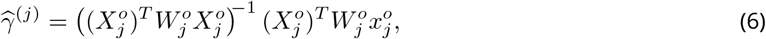

and there is no need for iteration.

The weights might be chosen in different ways. Here, we assume the noise term on measurements is additive, so at the same level for each measurement. In that case, one might apply

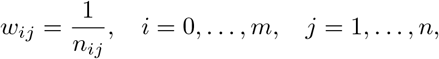

where *n_ij_* is the estimated background noise for that particular measurement. Within one replicate, *n_ij_* is roughly of the same size for all *i* = 0,…,*m*, but differs between replicates. Herein, we calculate the noise as explained in Section using the SuperSmoother method (Friedman, 1984).

We described here how to proceed when we have several replicates and we want to obtain an aggregated estimate of 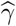. This will in turn produce an aggregated deamidation value for each peptide, as described in the next section. However, we keep separate estimates for each replicate by considering the special case when *n* =1 and replicate values are aggregated at a later step when the PQI is calculated (see Section 7).

#### Deamidation fractions

By normalisation

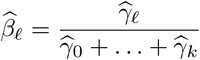

are estimates of the deamidation fractions corresponding to the parameters *β_ℓ_*, *ℓ* = 0,…, *k*, in Section. In our application of the theory we take *k* =1, and let *q* be the (least square estimated) fraction of the remaining, undeamidated Q, i.e. 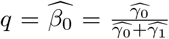. Herein, we define the fraction of undeamidated Q (*q*) as the extent of deamidation with *q* =1 meaning no deamidation whereas *q* = 0 referring to complete deamidation. Histograms of the extent of deamidation (*q*) of eight peptides are shown in Supplementary figure 1.

### Estimation of the overall deamidation level

We propose a linear mixed effect model, henceforth called the Parchment Glutamine Index (PQI) model, that integrates the estimated values of *q* and their analytical reliability with which the deamidation level is estimated. The Parchment Glutamine Index (PQI) model thus predicts an overall level of deamidation in a sample and an associated error of prediction.

#### PQI Model

The PQI model is a linear mixed effect model (LMM) that considers log-transformed q values as response variable with individual Peptide as the fixed effects, and Sample and Replicate as the random effects. As a result, the LMM fits the response variable at three different levels, namely, i) peptide, ii) sample, and iii) replicate. Herein, we use log-transformed q as the response variable to reflect the underlying kinetics of the loss of intact glutamine residues which follows pseudo-first order kinetics. Hence, the PQI model predicts the log-transformed deamidation level of a sample from the deamidation level of its individual peptides.

To simplify, we change the notation and structure of the data with respect to the previous section. Herein, the dataset is structured with log(*q*) values, *Reliability* estimates (see Section), factors that identifies Peptide (*P*, with *n_P_* levels), Sample (*S*, with *n_S_* levels) and Replicate (*R*, with *n_R_* levels). A summary of the dataset is given in Table 2.

**Table 2.**
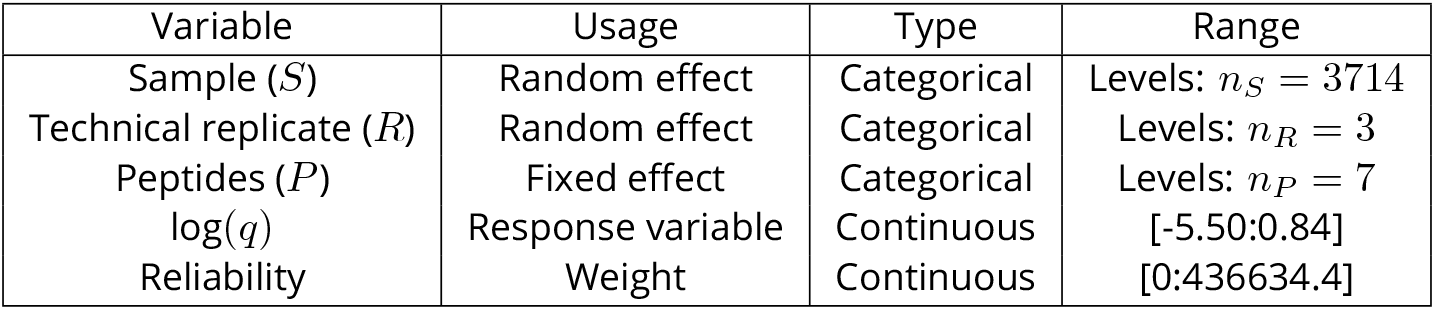
Summary of the dataset

Let *t* = 1,…,*u* denote the observation index, where *u* is the number of rows, and *u* = *n_S_*·*n_R_*·*n_P_*. The statistical model that we will use is the linear mixed effects model given by

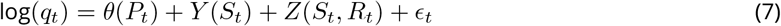

where,

- *θ*(*P*_1_),…, *θ*(*P_np_*) are the fixed effects of the each peptides,
- 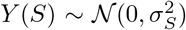 and 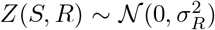 are the random components from sample, and replicate respectively, and
- 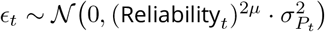 are the residuals.

The variance parameters for random effects are 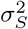 (sample, *S*) and 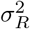 (replicate, *R*), and for the fixed effects of each peptide are 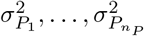. Furthermore, we scaled the residual variances by *Reliability* values generated from weighted square linear regression, see Eq. (5), to some power 2*μ*, and the PQI model estimates *μ*.

The aim is to predict *Y*(*s*) given the observations of log(*q_t_*) for indices t with *S_t_* = *s. Y*(*s*) is the random effect in the proposed linear mixed effect model that gives us the overall level of deamidation in a given sample, termed as PQI. To formalize this we define

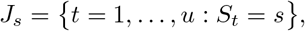

so that *J_s_* are the set of observation indices belonging to sample s. Additionally, we will use the following notation as given below:

- |*J_s_*| is the size of *J_s_*,
- *M*^T^ is the transpose of a matrix *M*,
- 1_*v*_ is the column vector of length *v* consisting of 1’s,
- *δ_x=y_* is the Dirac delta taking the value 1 when *x* = *y* and 0 otherwise,
- diag(*w*) is the diagonal matrix with the vector *w* in the diagonal.

Using this notation we have

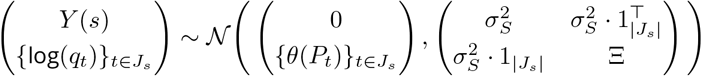

with 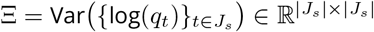 given by

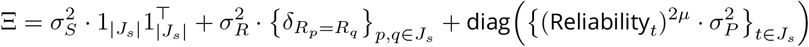

In particular, if we have observations of 3 replicates for all 7 peptides, then 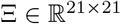 and it is given by

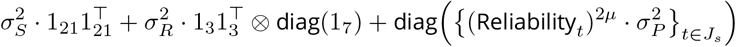

From the above joint normal distribution it follows by standard formulae that the conditional mean and the conditional variance of *Y*(*s*) are given by

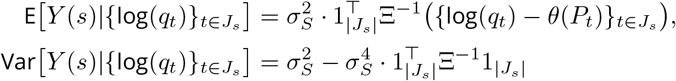

Note that E [*Y*(*s*)|{log(*q_t_*)}_*t*∈*J_s_*_ is the prediction of *Y*(*s*), and Var[*Y*(*s*) |{log(*q_t_*)}_*t*∈J_s__] is the associated prediction variance.

### Analysis workflow

We performed all the computations in the statistical programming language `R` (R Core Team, 2021) using the following packages: `nlme` (J Pinheiro et al., 2021), `dplyr` (Wickham et al., 2021), and `ggplot2` (Wickham, 2009). The prediction of *Y*(*s*) can be extracted directly from the `lme`-object using the function `nlme::ranef()`. However, the computation of the prediction variance requires implementation of the matrix formula. We developed an `R` package `MALDIpqi` for the whole workflow consisting of pre-processing of raw spectra, estimation of deamidation rates using weighted least squares linear regression, and applying linear mixed effect model to estimate the overall deamidation index of parchment (Figure 2), available in GitHub (https://github.com/ismaRP/MALDIpqi, https://doi.org/10.5281/zenodo.7105461).

**Figure 2.**
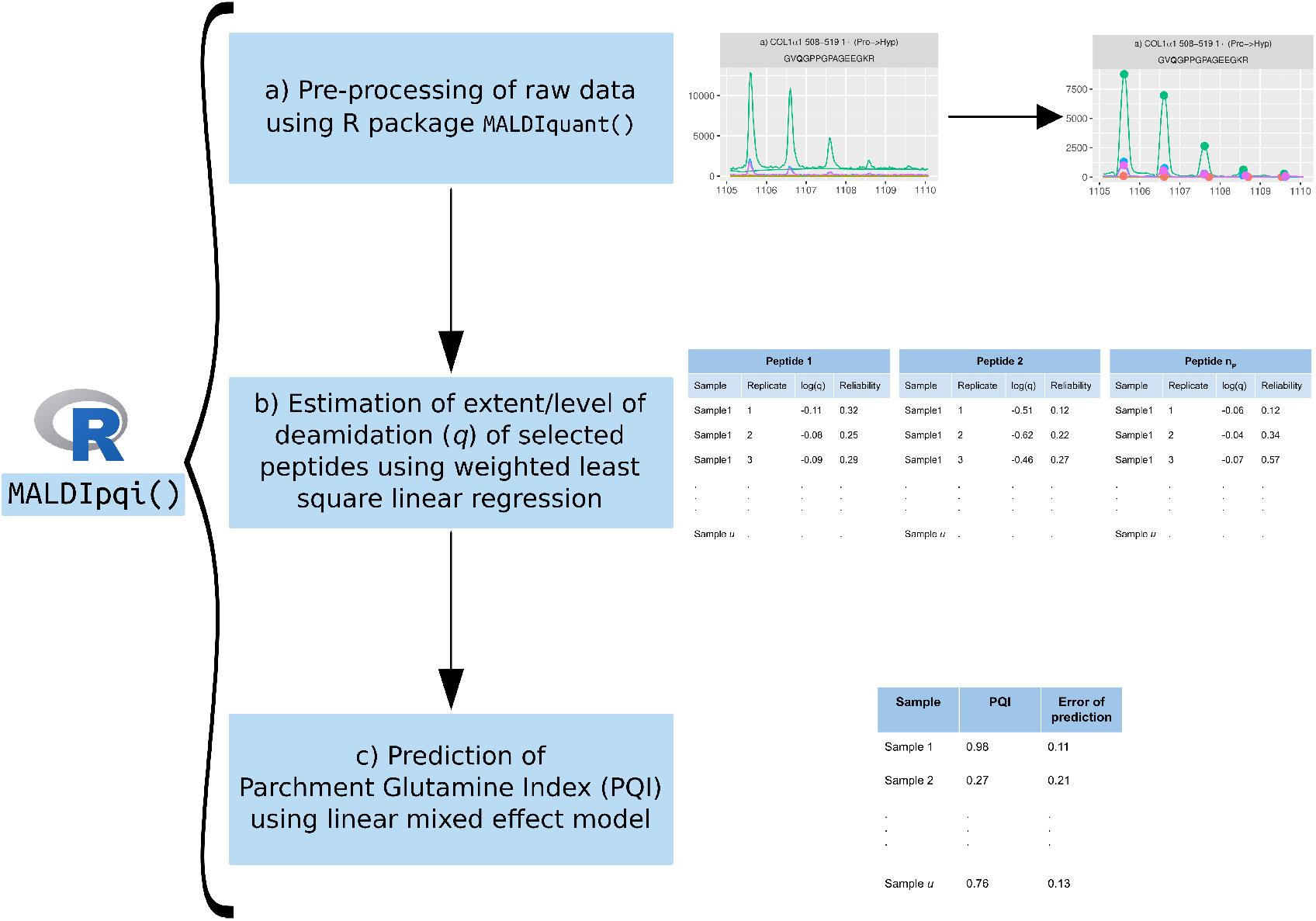
Summary of the workflow developed as an `R` package `MALDIpqi. MALDIpqi` consists of three steps, a) pre-processing of MALDI-TOF spectra, b) estimation of *q* of selected peptides, and c) prediction of PQI.

## Results and Discussion

We applied the workflow starting from pre-processing of raw data followed by estimation of deamidation levels of individual peptides and finally predicting the overall sample deamidation level, termed as PQI.

We let *q* denote the fraction of remaining non-deamidated Q (see Section) in the peptide under consideration. We estimated *q* values for selected eight peptides using weighted least squares linear regression on the isotopic distribution as obtained from MALDI-TOF spectra. Table 3 shows the first and third quartile of estimated q values to give an overview of deamidation levels in the peptides.

**Table 3.** Summary of the extent of deamidation in peptides and Restricted Maximum Likelihood estimates for fixed effects and relative rates of deamidation. Herein, *q* is the extent of deamidation in the peptides (from weighted least square linear regression) and 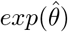 is the fixed effect estimates (from the PQI model).

| Peptide | 1st quartile of $q$ | Median $q$ | 3rd quartile of $q$ | $\exp(\hat{\theta})$ | Relative rates of deamidation |
| --- | --- | --- | --- | --- | --- |
| COL1α1 508-519 | 1.01 | 1.08 | 1.15 | 1.07 | 1.00 |
| COL1α1 270-291 | 0.93 | 1.06 | 1.14 | 1.03 | 6.56 |
| COL1α1 375-396 | 0.32 | 0.49 | 0.78 | 0.46 | 15.38 |
| COL1α1 934-963 | 0.82 | 1.00 | 1.18 | 0.98 | 5.71 |
| COL1α2 756-789 | 0.72 | 0.89 | 1.08 | 0.85 | 9.76 |
| COL1α2 756-789' | 0.64 | 0.78 | 0.93 | 0.74 | 10.34 |
| COL1α1 10-42 | 0.81 | 0.92 | 0.96 | 0.91 | 9.66 |
| COL1α1 10-42* | 0.63 | 0.79 | 0.94 | 0.74 | 9.41 |

### Relative rates of deamidation

Assuming the deamidation level over time follows first-order kinetics (NE Robinson, ZW Robinson, et al., 2004), then denoting the amount of non-deamidated Q of a particular peptide at time *t* by [*Q*]_*t*_, we have

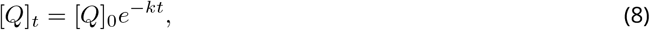

and hence

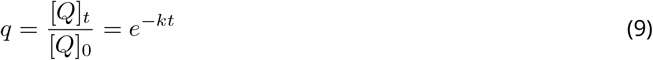

where, [*Q*]_0_ is the amount of Q at time 0, *k* is the deamidation rate constant, and *t* is the age of the sample.

Let *k*_1_ and *k*_2_ denote the deamidation rate constants of Peptide 1 and Peptide 2, respectively, from a particular sample. Similarly, let *q*_1_ and *q*_2_ denote the deamidation fractions of Peptide 1 and Peptide 2, respectively. Then the ratio of the deamidation rate constant of Peptide 2 to that of Peptide 1 can be expressed as

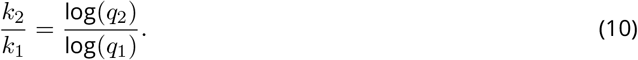

Considering Peptide 1 to deamidate slowly, we obtain a relative deamidation rate profile for each sample that might be compared across samples. Using this ratio overcomes the need to establish the absolute rate of deamidation. A limitation of this approach is when the levels of deamidation are very high, the true extent may be obscured due to the correspondingly high influence of noise in the spectra.

Among the eight peptides, the rates (see Table 3) are compared relative to COL1*α*1 508-519 (*m/z* 1105.58, VQG) which deamidates the most slowly. The only peptide which does not have a Glycine (Gly) C-terminal to the Gln is COL1*α*1 375-396 (*m/z* 2040.97, GQD), and this is the most rapidly deamidated. This rapid deamidation explains the clustering of fitted values towards the left of the residual plot for this peptide, as shown in Figure 3. COL1*α*1 934-963 (*m/z* 2689.25), contains two glutamine residues both oriented in the same plane (PQGFQG), but even their combined rate is nevertheless slower than COL1*α*1 375-396. Curiously *m/z* 3084.42 (identified by Mascot (Perkins et al., 1999) as COL1*α*1 10-42) has a rate of deamidation which is one third that of *m/z* 3116.40, which was interpreted as the same peptide but with only two less oxygen atoms. The most probable explanation is that one of these peptides may have been misidentified, as it seems unlikely that additional oxidation/hydroxylations would have such a significant effect on the rate of deamidation.

**Figure 3.**
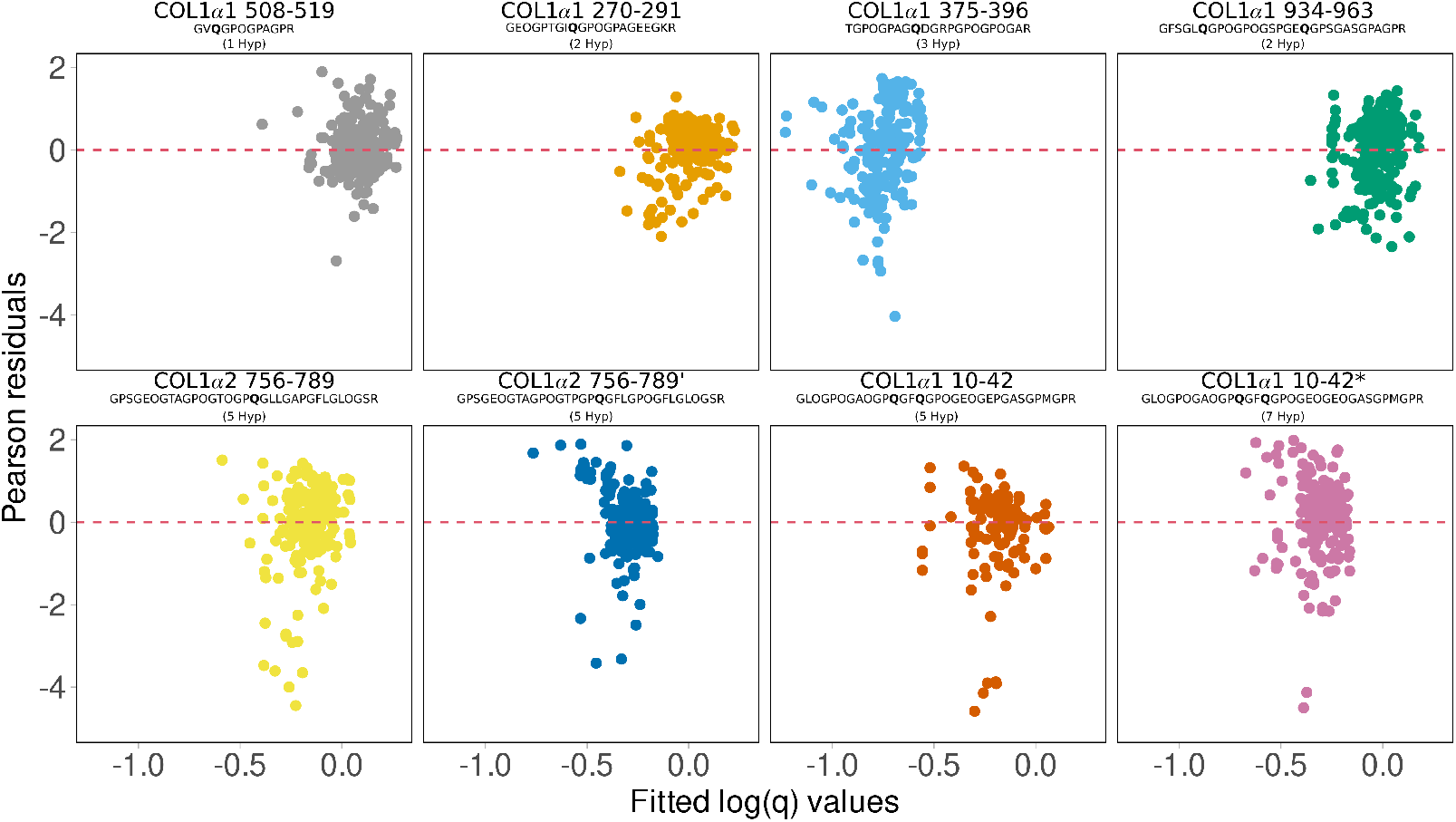
Residual plots between fitted values and Pearson residuals for a) COL1*α*1 508-519, b) COL1*α*1 270-291, c) COL1*α*1 375-396, d) COL1*α*1 934-963, e) COL1α2 756-789, f) COL1α2 756-789’, g) COL1*α*1 10-42 (5 Pro →Hyp), and h) COL1*α*1 10-42* (7 Pro →Hyp), explores the model fitting quality. Random distribution of standardised residuals around 0 within ±2 suggests that the proposed linear mixed effect model fits well. However, there are a few badly modelled q values for some of the peptides. 200 randomly selected data points are shown in each plot.

A recent study (Simpson et al., 2019) demonstrated that MALDI technique underestimates peptides with E in peptide mixtures containing Q and E substitutions that could potentially result in the systematic overestimation of q, i.e. remaining Q. Since we are considering relative rate differences between samples, this effect does not play a crucial role in our analyses.

The log-transformed q values were then transferred to the PQI model, which fits the deamidation at peptide level and predicts the sample level deamidation. We used the lme function in R from the package nlme to fit the linear mixed effect model using restricted maximum likelihood (REML). PQI model estimates for sample level variance 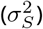 is 0.01 and replicate level variance 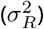 is 4.09 * 10^-11^ with *μ* = −0.06. The back-transformed peptide level fixed effect estimates and the relative levels of deamidation are given in Table 3.

We validated the model fitting using residual plots (residuals vs. fitted values) and normal quantile-quantile plots of the Pearson residuals (residuals standardized by their estimated standard deviation). Residual plots are the most common diagnostic tool to assess the constant variation of residuals (JC Pinheiro and Bates, 2000). Diagnostic plots for PQI model fitting for the slowest deamidating peptide, COL1*α*1 508-519, are shown in Figure 4 showing a valid statistical model except slightly too heavy tails in the normal distribution. Pearson residuals for almost all samples are randomly distributed around 0 with magnitudes ranging between ±2, and without any concerning patterns as depicted in Figure 3.

**Figure 4.**
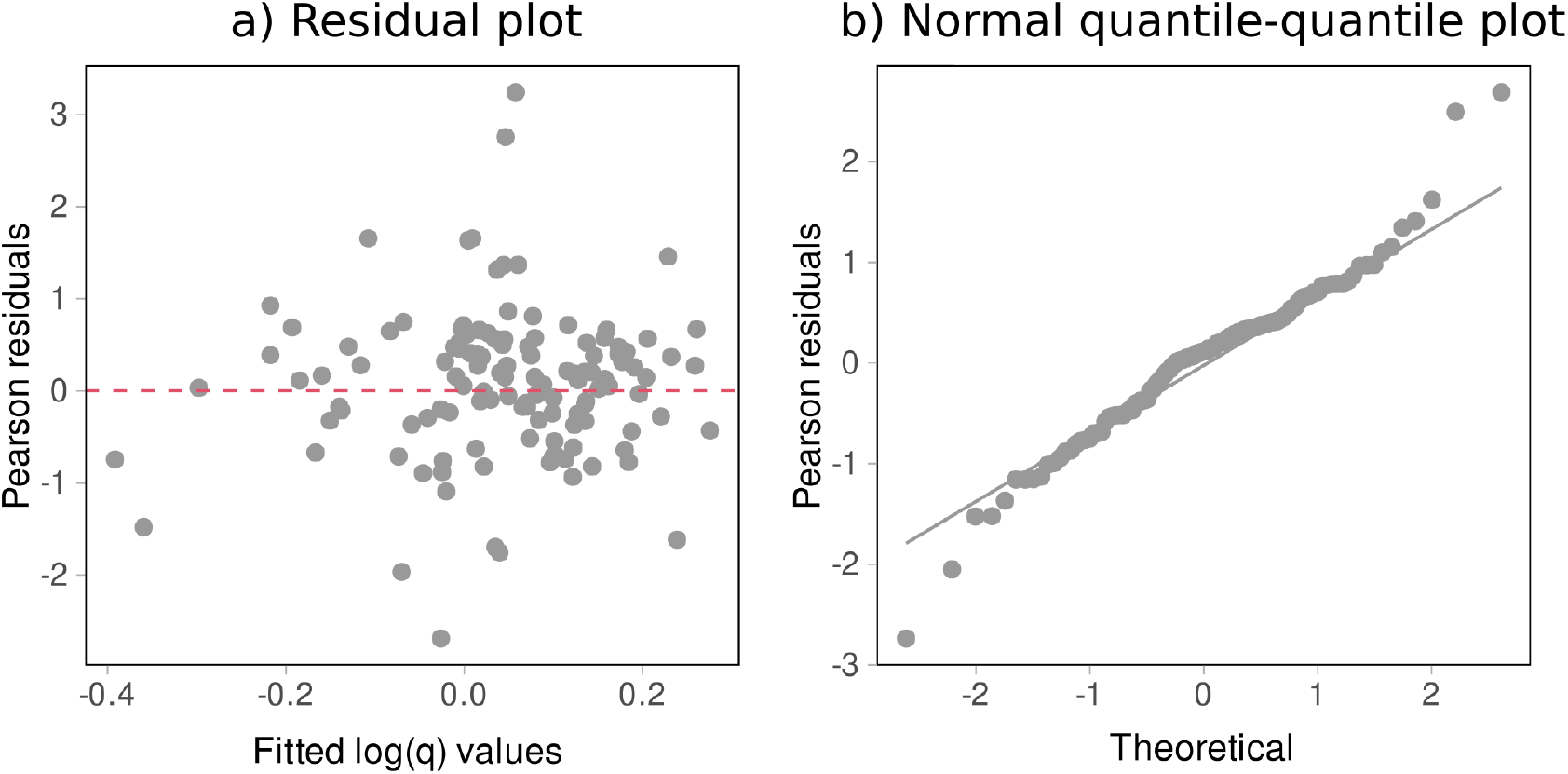
Diagnostic plots for PQI model fitting for COL1*α*1 508-519 indicating a valid statistical model. a) Residual plot between fitted log(*q*) values and Pearson residuals, and b) quantile-quantile plot of residuals wherein the grey line indicates the 1-1 line. Both diagnostic plots indicates that the PQI model is valid except slightly too heavy tails in the normal distribution. 200 randomly selected data points are shown in each plot.

Normal quantile-quantile plots compare quantiles of Pearson residuals to quantiles of standard normal distribution. Linearity of the quantile-quantile plot implies that residuals are normally distributed as proposed in the Parchment Glutamine Index (PQI) model. With the exception of a few data points on both tails of the quantile-quantile plots, the model fits the deamidation well (Figure 5). The few data points that do not fall onto the quantile-quantile line for peptides COL1α2 756-789 and COL1α2 756-789’ (see Figure 5) is the result of a low S/N that affects the correct estimation of *q* values from the MALDI-TOF spectra.

**Figure 5.**
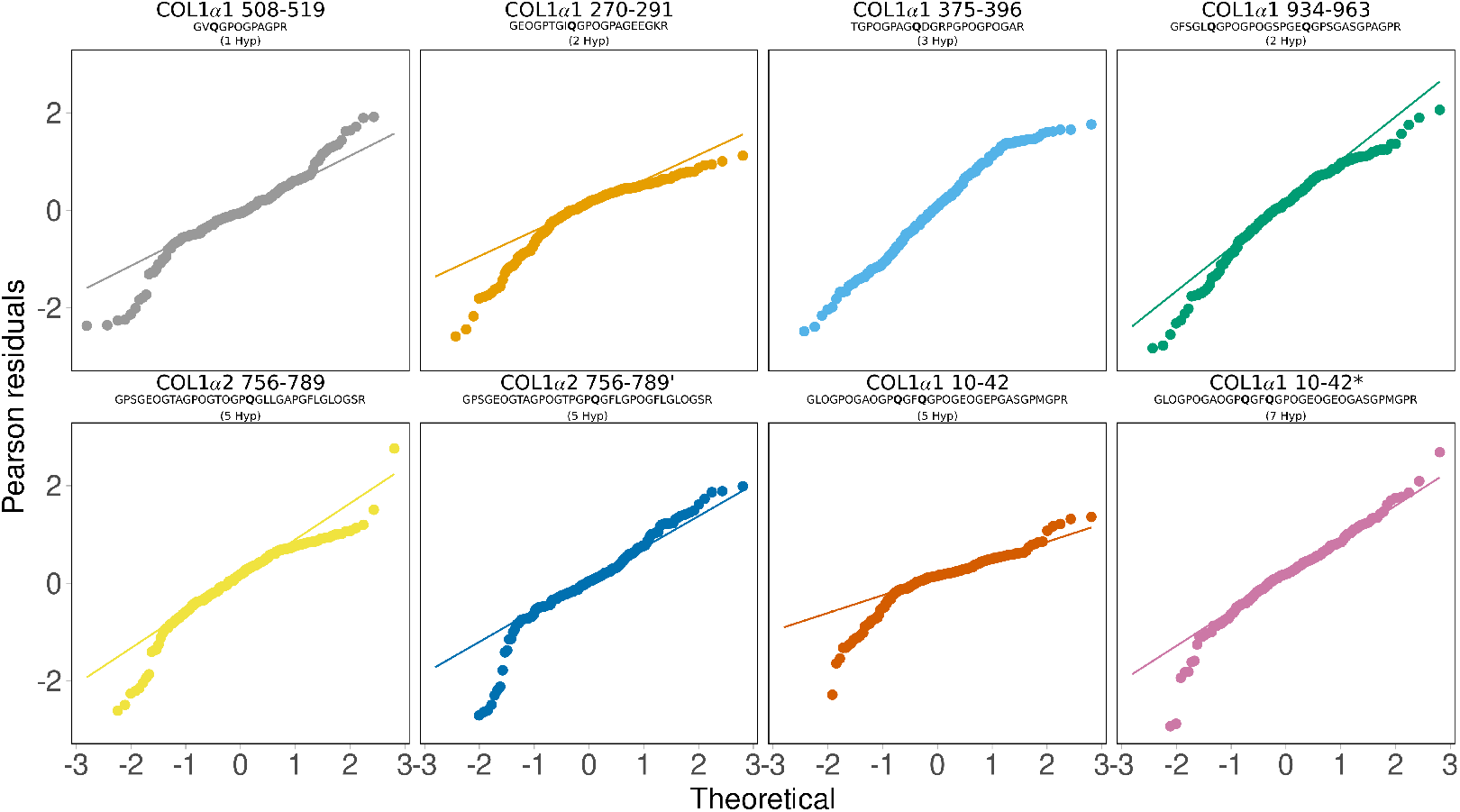
Normal quantile-quantile plots of Pearson residuals for the model fit for a) COL1*α*1 508-519, b) COL1*α*1 270-291, c) COL1*α*1 375-396, d) COL1*α*1 934-963, e) COL1α2 756-789, f) COL1α2 756-789’, g) COL1*α*1 10-42 (5 Pro →Hyp), and h) COL1*α*1 10-42* (7 Pro →Hyp). Except a few deviations from the inserted 1-1 lines, in particularly at the tails for some of the peptides, the quantile-quantile plots indicates normality of the residuals as proposed by the PQI model. 200 randomly selected data points are shown in each plot.

The PQI model predicts the sample level log(*q*) value and we therefore argue that the exp(log(*q*)) value depicts the overall extent of deamidation in a sample, termed as the Parchment Glutamine Index (PQI). From the samples considered in the analysis, PQI predicted from the model ranges from 0.47 to 1.26 with 54% of the values above 1, although theoretically the full PQI range is from 0 to 1. A low value of PQI implies more liming and hence low quality of parchment while a value of 1 indicates no deamidation. The model generates some PQI values greater than 1 which we attribute to the problem of accurate baseline correction and noise. We note that replicates generally produce concordant deamidation estimates and different peptides from the same sample generally produce highly correlated PQI values. It is worth noting that (Wilson et al., 2012) reported a similar problem and used truncation to solve for deamidation fractions > 1. A histogram of predicted PQI values are shown in Supplementary figure 2.

From the PQI model, we estimated peptide level fixed effects and sample level random effects. Whilst the fixed effect is the mean log(*q*) of each peptide, the random effect *Y*(*s*) is the predicted overall deamidation level in a sample, PQI (see Section). A few q values for the peptides COL1α2 756-789 and COL1α2 756-789’were not fitted well in the PQI model implying inaccurate estimates from spectral peaks with low S/N. Relative to COL1*α*1 508-519, COL1*α*1 375-396 displayed higher rates of deamidation where as COL1*α*1 270-291 displayed lowest rate (see Table 3).

### Applications of PQI

As an illustration of the application of the PQI model we explore levels of deamidation from a collection of manuscripts from the library at Orval Abbey, Belgium (Ruffini-Ronzani et al., 2021) by comparing PQI values with species, thickness, typology, and production period as shown in Figure 6. We expect no link between deamidation and time or preservation histories of parchment. This is due to the fact that time-dependent deamidation via hydrolysis is a much slower mechanism than the deamidation produced by the aggressive liming of animal skins.

**Figure 6.**
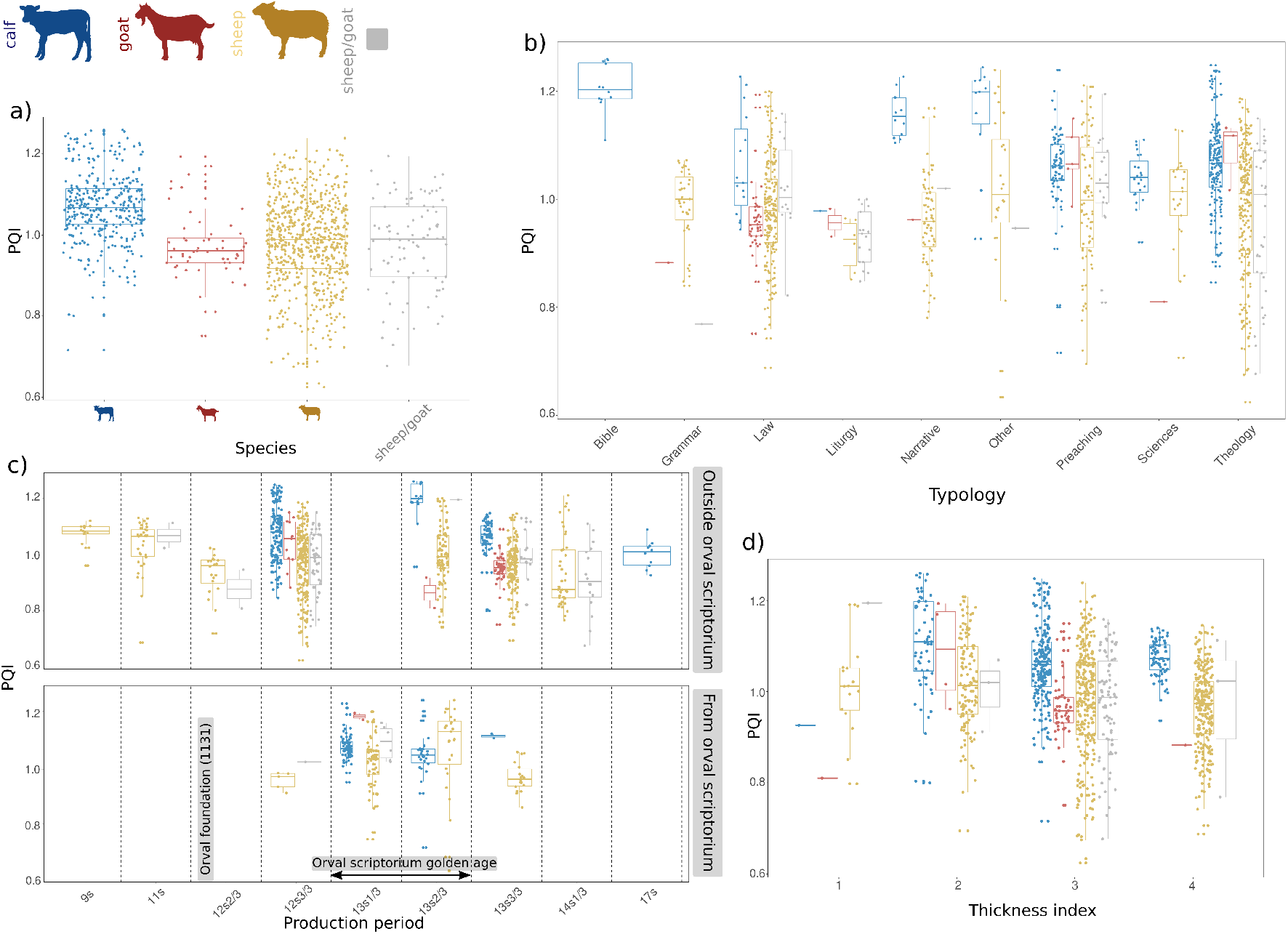
Plots depicting applications of PQI: a) Comparison of PQI against different species used for the production of parchment depicting that manuscripts made out of calfskin were of better quality than the ones made with sheepskin or goatskin. b) Comparison of PQI against different typology the parchments were used for. Biblical manuscripts were written on calfskin, having the highest PQI, which is in accordance with the findings in (Ruffini-Ronzani et al., 2021). Sheepskin was commonly used to produce grammar and theology texts with an intermediate deamidation index. c) Comparison of PQI against production period for parchment locally produced in Orval scriptorium (bottom panel) and for imported parchments (top panel) starting from 9th century until 17th century. The timeline is organised by thirds of a century (early, mid, and late). Orval scriptorium was founded in the early 12th century. The use of calfskin to produce parchments remained constant during the “golden age”(first third and second third of the 13th century) of the scriptorium. d) Comparison of PQI against the thickness indices of codicological units. The thickness was determined depending on the number of folios in the codicological unit (Thickness index; 1 = less than 10 folia, 2 = 11-100 folia, 3 = 101-200 folia, 4 = greater than 200 folia.). (Icons of calf, goat, and sheep created with BioRender.com)

#### PQI vs species

PQI varies with species, parchment produced from calf has higher PQI values, than those produced from sheep or goat (Figure 6a). Goat skin parchments had the lowest PQI, suggesting that they were the most aggressively limed. We also observe the highest PQI values in calfskin used for Bible, and we speculate that this would probably have been perceived as of the best quality. Law and science texts tended to use the lowest quality parchment, although within each group of texts there was considerable variation (Figure 6b).

#### PQI vs parchment thickness

Sheep showed the widest range of values, and goat had the lowest PQI values (most deamidated). Estimated thickness suggests that the (small number of) very finest parchment (Thickness index 1) are not of the best quality, an unexpected finding which should be explored further. There is nevertheless a gradual fall in PQI in the next three thickness groups as might be expected, with the greatest levels of deamidation in the coarsest membranes as shown in Figure 6d.

#### PQI vs time

A temporal comparison of parchment production from the 9th century until the 17th century reveals the highest PQI values occurred during the “golden age” of the Orval scriptorium (first half of the 13th century), presumably before the disastrous fire of 1252 (see Figure 6c).

## Conclusion

The PQI model allows us to reliably estimate the quality of parchment production by deriving an index which combines the extent of deamidation of seven tryptic peptide markers from MALDI-TOF analysis, applied in bioarchaeology, termed as ZooMS for species identification. MALDI-TOF is the widely used proteomics based method in bioarchaeology, as it is the basis of ZooMS, due to its reduced costs and easiness of sample processing. Although one can obtain accurate PQI estimates from high-quality data generated using high-resolution mass spectrometers, our workflow is able to maximize the information obtained from existing data generated from ZooMS studies. It uses a three step workflow, the pre-processing of spectra for optimal assessment of each mass envelope, estimating deamidation levels in peptides using weighted least square linear regression, and finally, predicting the overall deamidation level in a sample using a linear mixed effect model. Each step is coded in R as a package `MALDIpqi()`, enabling high throughput analysis of large datasets.

We applied the workflow to 3714 MALDI-TOF spectra from parchments in the library of the Orval Abbey and were able to observe a number of patterns. There is a large variation in PQI between membranes but some patterns are evident. Coarser membranes are more heavily limed than thinner folia, and calfskin is more gently processed than sheep and goatskin. Both of these would be anticipated based upon our knowledge of parchment production, although we were surprised by the low PQI values of goatskin, which is typically less fatty than sheepskin and therefore does not require such long exposure to saponify and hence remove lipids. More subtle observations are also apparent at Orval Abbey; texts acquired after the fire of 1252 are on average worse than those acquired during the so-called golden age which preceded it.

In addition to this biocodicological application of PQI, livestock collagen is widely used in the food industry and biomedicine. Therefore the developed three step workflow offers a simple method to assess levels of Gln deamidation of processed collagen.

## Acknowledgements

We thank Julie Wilson (JW) for helpful comments and advice. Preprint version 6 of this article has been peer-reviewed and recommended by Peer Community In Archaeology of the PCI (https://doi.org/10.24072/pci.archaeo.100019).

## Data, script and code availability

Data are available online: https://doi.org/10.5281/zenodo.5648106

Script and codes are available online: https://doi.org/10.5281/zenodo.7105461

## Supplementary information availability

Supplementary information is available online: https://doi.org/10.24072/pci.archaeo.100019

