## Supplementary material for "Parchment Glutamine Index (PQI): A novel method to estimate glutamine deamidation levels in parchment collagen obtained from low-quality MALDI-TOF data": Supplementary material.pdf

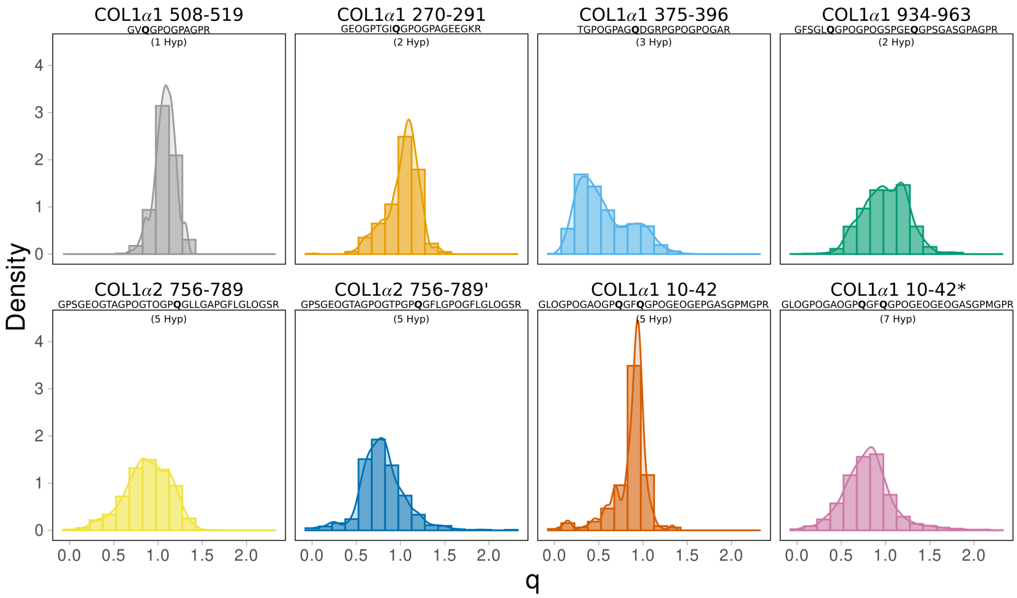


**Supplementary figure 1: Histogram of extent of deamidation ($q$) for the 8 peptides.**

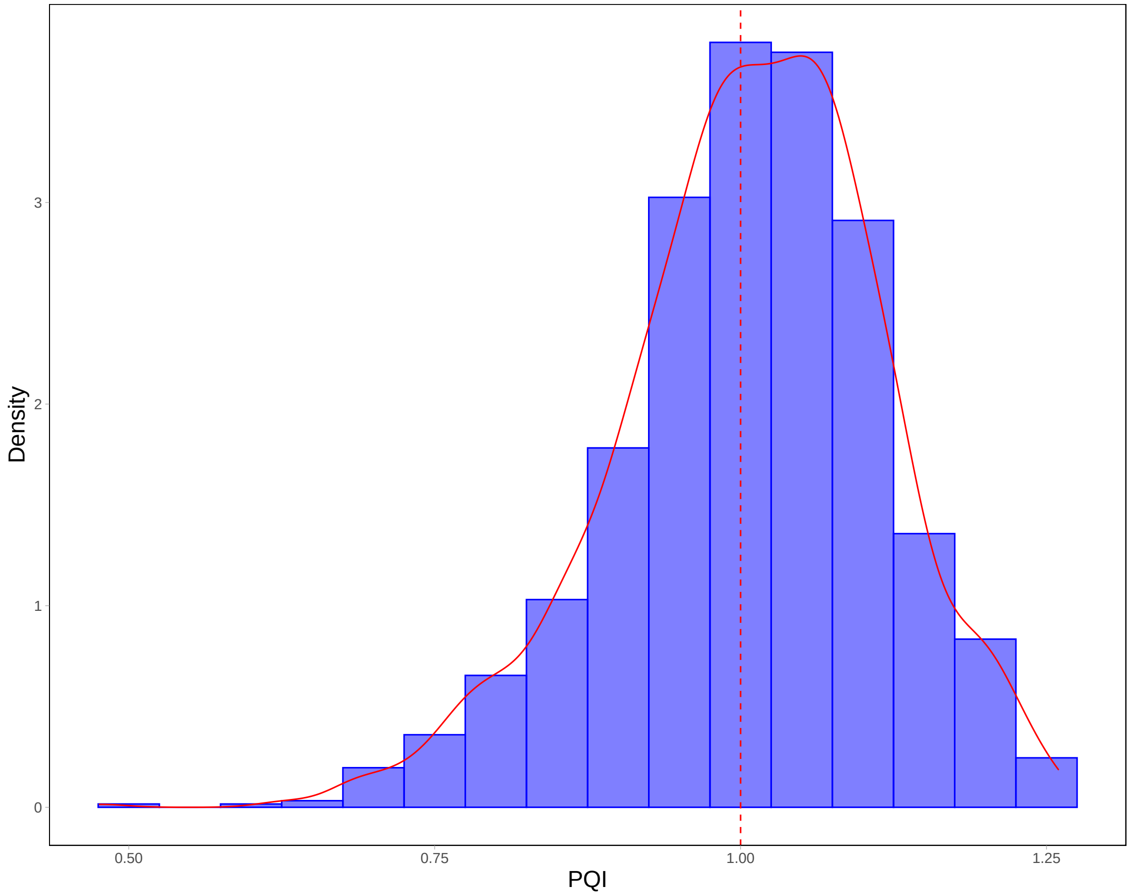


**Supplementary figure 2: Histogram of Parchment Glutamine Index (PQI).** Although theoretically the value of PQI ranges from zero to one, 54% of the values are above one due to the problem of accurate baseline correction.
